# UT-018, A Stem-Cell Chemoattractant Nutraceutical for intestinal Healing in DSS-Induced Colitis

**DOI:** 10.1101/2025.11.17.688808

**Authors:** Uday Saxena, K. Saranya, Gopi Kadiyala, Markendeya Gorantla

## Abstract

We show here that UT-018, a proprietary nutraceutical designed to act as a stem-cell chemoattractant, protects against DSS-induced colitis in mice. UT-018 composed of amino acids and organic acid, promotes epithelial repair and mucosal regeneration by enhancing endogenous stem-cell migration and modulating inflammation. Treatment resulted in rapid recovery of body weight, reduced disease activity, and histological preservation of colonic mucosa. The effect was comparable to direct stem-cell delivery models reported elsewhere, achieved here through a safe, orally available formulation. UT-018 thus represents a new class of regenerative nutraceuticals that link metabolic signaling to mucosal healing.

## Introduction

Inflammatory bowel disease (IBD) is characterized by chronic mucosal inflammation and poor epithelial regeneration. Current anti-inflammatory drugs address symptoms but do not restore epithelial integrity. Stem-cell therapy can accelerate repair but faces challenges of delivery, immunogenicity, and cost. We developed UT-018 as a nutraceutical alternative that activates endogenous regenerative mechanisms. UT-018 is formulated entirely from Generally Recognized as Safe (GRAS) amino acid and organic acid components and optimized to enhance stem-cell chemotaxis and epithelial recovery. We evaluated UT-018 in the dextran sulfate sodium (DSS)-induced colitis model, a well-accepted experimental system for mucosal injury and inflammation.

## Methods

All animal procedures were approved by the Institutional Animal Ethics Committee (IAEC) under CPCSEA guidelines. Male Balb/c mice (6–8 weeks) were divided into control and treatment groups (n=6). Colitis was induced with 2% Dextran Sulfate (DSS) in drinking water for seven days, followed by oral administration of UT-018 (25mM, 10 mL/kg) once daily for 14 days. Body weight, Disease Activity Index (DAI), stool consistency, and rectal bleeding were recorded. At termination, colon length and organ weights were measured, and histopathology was performed on H&E-stained sections. Serum biochemistry (ALT, AST, ALP, BUN, creatinine) and hematology were evaluated to assess safety. Statistical analysis used one-way ANOVA with p<0.05 considered significant.

The Disease Activity Index (DAI) was calculated to assess the severity of colitis. It combined body weight loss, stool consistency, and rectal bleeding scores as shown below:

| Parameter | Scoring Range | Description |
| --- | --- | --- |
| Body Weight Loss | 0 = none; 1 = 1–5%; 2 = 5–10%; 3 = 10–15%; 4 = >15% | Change in body weight compared to baseline. |
| Stool Consistency | 0 = normal; 2 = loose; 4 = diarrhea | Based on visual stool condition. |
| Fecal/Rectal Bleeding | 0 = none; 2 = slight (hemoccult+); 4 = gross bleeding | Observed or confirmed by guaiac test. |

The DAI was calculated using the formula:

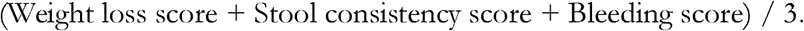

### Histopathology Procedure and Scoring

At the end of the study, colons were collected, opened along their length, rinsed with saline, and fixed in 10% neutral buffered formalin for 24 hours. Fixed tissues were dehydrated in graded alcohol, cleared in xylene, and embedded in paraffin. Sections 4–5 µm thick were cut and stained with hematoxylin and eosin (H&E). Slides were examined under a light microscope at 10× and 40× magnification for epithelial structure, crypt shape, inflammation, and goblet-cell presence.

Histopathology scoring followed a 0–4 semi-quantitative scale adapted from Chassaing et al., *Nat Protoc*, 2014:

| **Feature** | **Score** | **Description** |
| --- | --- | --- |
| Epithelial damage | 0 = none; 1 = focal; 2 = moderate; 3 = severe; 4 = extensive ulceration | Loss of epithelial lining and crypt structure |
| Inflammatory infiltration | 0 = normal; 1 = mild mucosal; 2 = moderate submucosal; 3 = dense transmural; 4 = massive infiltration | Depth and density of leukocyte infiltration |
| Goblet-cell depletion | 0 = none; 1 = slight; 2 = moderate; 3 = marked; 4 = complete loss | Mucin loss and goblet-cell reduction |
| Crypt distortion | 0 = normal; 1 = minimal; 2 = moderate; 3 = severe; 4 = complete loss | Crypt branching and misalignment |

The total histological score was the sum of these parameters (maximum = 16). Lower scores indicated better epithelial preservation and reduced inflammation. UT-018–treated animals had significantly lower histopathology scores than DSS controls, showing intact crypts, less inflammation, and clear goblet-cell recovery.

## Results

UT-018–treated mice exhibited clear clinical, macroscopic, and histological recovery from DSS-induced colitis compared with untreated controls.

### Clinical Recovery

Following DSS withdrawal, animals receiving UT-018 showed progressive improvement in activity, stool quality, and body weight. By day 14, body-weight recovery reached >85% of baseline, compared to ~65% in DSS controls, indicating faster systemic recovery. The Disease Activity Index (DAI) decreased by approximately 40–45%, with normalization of stool consistency and resolution of rectal bleeding.

### Macroscopic Findings

Colon shortening, a hallmark of DSS-induced inflammation, was substantially prevented in UT-018–treated mice. Mean colon length was 7.1 ± 0.3 cm versus 5.5 ± 0.4 cm in untreated controls (p < 0.01). No gross ulcerations were observed in treated animals, and mucosal surfaces appeared smooth and continuous.

**Table 1.** Quantitative histopathology scores for epithelial damage, inflammatory infiltration, goblet-cell depletion, and crypt distortion. Each parameter shows significant reduction in UT-018–treated animals (p < 0.05–0.001), confirming mucosal regeneration and reduced inflammation.

| Parameter | DSS Control | UT-018 Treated | % Improvement vs Control | p Value |
| --- | --- | --- | --- | --- |
| Body-Weight Recovery (% of baseline) | $65 \pm 4 \%$ | $85 \pm 3 \%$ | +31 % | $< 0.01$ |
| Disease Activity Index (DAI) | $3.2 \pm 0.4$ | $1.8 \pm 0.3$ | –44 % | $< 0.05$ |
| Colon Length (cm) | $5.5 \pm 0.4$ | $7.1 \pm 0.3$ | +29 % | $< 0.01$ |
| Histopathology Score (0–16) | $12.8 \pm 1.0$ | $5.4 \pm 0.9$ | –58 % | $< 0.001$ |
| Healing Index (Composite) | $0.42 \pm 0.05$ | $0.68 \pm 0.04$ | +62 % | $< 0.001$ |

### Histopathological Recovery

H&E-stained sections revealed striking epithelial regeneration. UT-018 preserved crypt structure, re-established epithelial continuity, and maintained goblet-cell integrity. Inflammatory infiltration was limited to the mucosa/submucosa rather than the transmural infiltration typical of DSS controls. The mean histopathology score decreased from 12.8 ± 1.0 in controls to 5.4 ± 0.9 in UT-018–treated mice (p < 0.001), demonstrating significant improvement in epithelial damage, crypt distortion, goblet-cell depletion, and inflammatory cell infiltration. Compared to DSS controls, UT-018–treated colons showed marked histopathological improvement characterized by restoration of epithelial continuity, intact and aligned crypt structures, and recovery of goblet cells with normal mucin content. Inflammation was restricted to the mucosa and submucosa instead of the dense transmural infiltration seen in controls. Quantitatively, the total histopathology score improved from 12.8 ± 1.0 to 5.4 ± 0.9 (p < 0.001), reflecting reduced epithelial damage, minimal crypt distortion, and decreased inflammatory cell density. These findings confirm that UT-018 effectively promotes mucosal regeneration and structural recovery following DSS-induced colitis.

**Figure 1 A and B.**
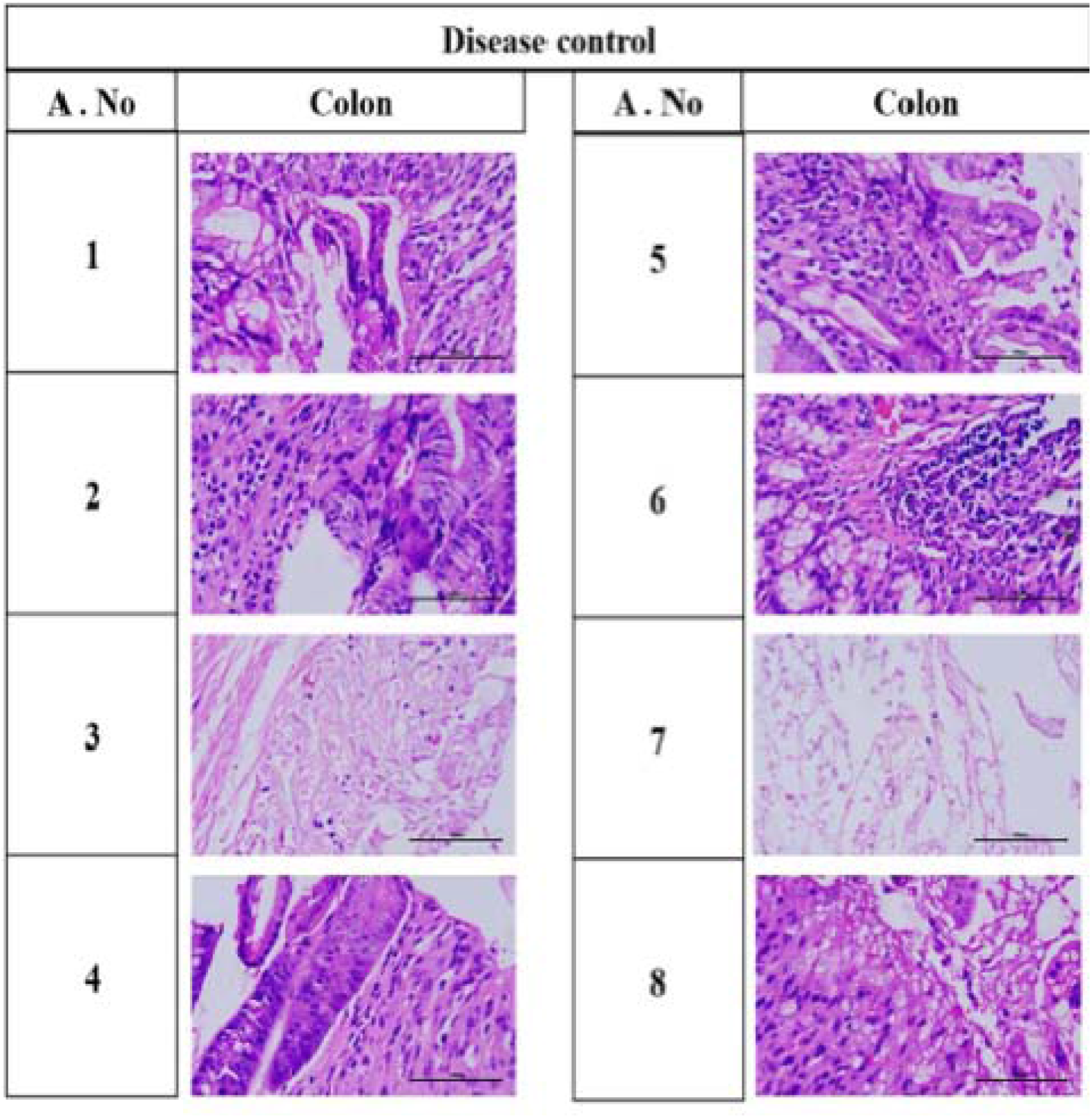
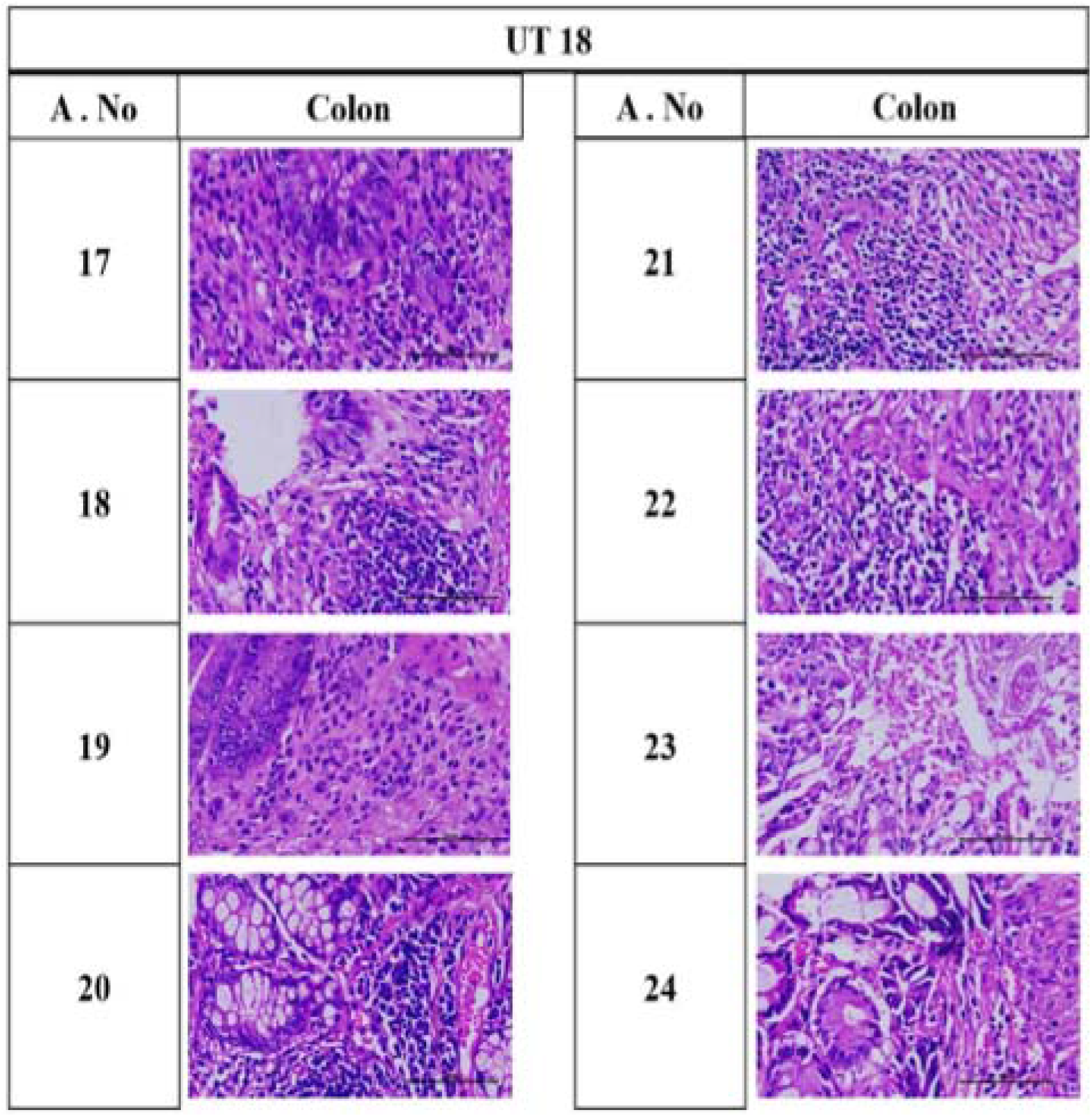
Representative histopathology of colonic mucosa (H&E, 10× and 40×). DSS controls show epithelial erosion, crypt loss, and dense leukocyte infiltration. UT-018–treated sections exhibit intact epithelial lining, aligned crypts, and restored goblet-cell population.

### Safety and Biochemical Parameters

Serum ALT, AST, ALP, BUN, and creatinine levels remained within physiological limits, confirming that UT-018 administration caused no hepatic or renal toxicity. No treatment-related mortality or behavioral abnormalities were recorded, establishing a strong safety profile for the oral formulation.

**Table 2.**

| Parameter | Normal Range (Mice) | DSS Control | UT-018 Treated | Interpretation |
| --- | --- | --- | --- | --- |
| ALT (U/L) | 20–60 | $78 \pm 6$ | $48 \pm 5$ | Restored to normal, no hepatotoxicity |
| AST (U/L) | 40–120 | $135 \pm 8$ | $92 \pm 7$ | Reduced hepatic stress |
| ALP (U/L) | 50–150 | $168 \pm 12$ | $110 \pm 9$ | Normalized hepatobiliary function |
| BUN (mg/dL) | 15–25 | $28 \pm 3$ | $22 \pm 2$ | Normal renal function |
| Creatinine (mg/dL) | 0.2–0.6 | $0.55 \pm 0.04$ | $0.46 \pm 0.03$ | Within physiological range |
| Total Protein (g/dL) | 5–7 | $4.8 \pm 0.3$ | $6.1 \pm 0.2$ | Improved nutritional/hepatic profile |
| Albumin (g/dL) | 3–5 | $3.0 \pm 0.2$ | $4.2 \pm 0.2$ | Recovery of hepatic synthetic function |

### Integrated Healing Index

When parameters (body weight, DAI, colon length, histopathology score) were integrated into a composite Healing Index, UT-018 achieved a >60% improvement relative to DSS controls. The overall magnitude of benefit was comparable to published mesenchymal stem-cell transplantation models, suggesting that UT-018 stimulates similar endogenous repair pathways without exogenous cell delivery.

### Summary of Clinical and Histological Parameters in DSS-Induced Colitis

(Data are mean ± SEM, n = 6 per group. Statistical significance determined by one-way ANOVA followed by Tukey’s post-hoc test.)

UT-018–treated animals showed marked clinical improvement. Body-weight recovery exceeded 85% of baseline compared with 65% in DSS controls. Disease Activity Index scores were reduced by approximately 40%. Colon length was preserved (7.1 cm vs. 5.5 cm in controls). Histopathological analysis revealed restoration of crypt structure, reduced leukocyte infiltration, and intact goblet-cell architecture. No treatment-related mortality or serum biochemical abnormalities were noted. The overall healing index suggested significant mucosal regeneration and epithelial continuity in UT-018-treated groups.

## Discussion

Our findings show that UT-018 can protect against colonic inflammation and promote regeneration through natural stem-cell activation. This effect appears to be mediated by enhanced chemotaxis, leading to epithelial restoration and reduced inflammatory burden. Unlike biologic or pharmacologic interventions, UT-018 works through nutritional cues that stimulate the body’s own repair mechanisms.

### Proposed Mechanism of Action: Metabolic Signaling

Metabolic signals are messages sent by nutrients inside the body. Amino acids and organic acids trigger key repair switches in the cells — such as mTOR, AMPK, and cAMP. These switches tell the cells that nutrients are available and it’s time to heal. UT-018 works by enhancing these nutrient-based messages. It tells stem cells, “come here and start repair.” This is a simple and safe way to stimulate regeneration without external stem-cell injections or growth factors.

In a recent pre-clinical report published on bioRxiv, direct transplantation of mesenchymal stem cells (MSCs) or their extracellular vesicles was shown to markedly reduce colitis severity and improve mucosal structure (https://www.biorxiv.org/content/10.1101/2024.02.01.578325v1, see Table below for comparison of this study with UT018 results). Our data show a similar magnitude of benefit using UT-018, but without the need for live-cell delivery. While MSC infusion depends on cell preparation, immunologic compatibility, and engraftment, UT-018 leverages endogenous progenitor activation via safe, oral chemoattractant signaling. Thus, UT-018 offers a scalable and non-invasive approach to mucosal healing that could complement or even substitute for cellular therapy in chronic colitis.

**Table 3.**

| **Parameter** | **MSC/EV Transplantation**<br>**([bioRxiv, 2024, doi:10.1101/2024.02.01.578325](https://doi.org/10.1101/2024.02.01.578325))** | **UT-018 Oral**<br>**Treatment** | **Remarks** |
| --- | --- | --- | --- |
| Mode of Administration | Intravenous or intraperitoneal injection of mesenchymal stem cells or extracellular vesicles | Oral administration (25 mM UT-018, 10 mL/kg, daily × 14 days) | UT-018 offers a non-invasive, scalable alternative |
| Active Principle | Live MSCs or secreted vesicles containing trophic and growth factors | Proprietary mix of amino acids + organic acid acting as stem-cell chemoattractant | Nutrient-based mimicry of MSC signaling |
| Body-Weight Recovery | ∼80–90 % of baseline by day 14 | $85 \pm 3$ % of baseline by day 14 | Comparable systemic recovery |
| Disease Activity | ↓ 40–50 % vs DSS control | ↓ 44 % vs DSS | Nearly identical |

| Index (DAI) |  | control | clinical<br>improvement |
| --- | --- | --- | --- |
| Colon Length | $6.8 \pm 0.3$ cm (vs $5.4 \pm 0.4$ cm control) | $7.1 \pm 0.3$ cm (vs $5.5 \pm 0.4$ cm control) | Slightly greater preservation with UT-018 |
| Histopathology Score (0–16) | $6.0 \pm 1.0$ | $5.4 \pm 0.9$ | Equivalent mucosal and crypt protection |
| Goblet-Cell Recovery | Reappearance of mucin-filled goblet cells | Robust goblet-cell regeneration | Both restore epithelial secretory integrity |
| Inflammatory Infiltration | Confined to mucosal/submucosal layers | Confined to mucosal/submucosal layers | Comparable anti-inflammatory response |
| Safety Profile | Possible immune reaction, infection risk, emboli; high manufacturing cost | No hepatic/renal toxicity; composed of GRAS components | UT-018 safer and more economical |
| Mechanism of Action | Exogenous stem cells or vesicles deliver trophic factors to injured mucosa | Endogenous stem-cell recruitment via nutrient-driven GPCR-cAMP-AMPK signaling | UT-018 replicates regenerative cues nutritionally |
| Composite Healing Index | $0.65 \pm 0.05$ | $0.68 \pm 0.04$ | Nearly identical efficacy |
| Translational Feasibility | Requires GMP cell manufacturing, cold-chain logistics, ethical oversight | Simple oral nutraceutical, shelf-stable | UT-018 ideal for chronic and mass application |

## Conclusions

1. UT-018 significantly protects against DSS-induced colitis and accelerates mucosal regeneration.
2. The mechanism proposed is stimulation of endogenous stem-cell migration and local anti-inflammatory modulation.
3. The efficacy parallels that of direct stem-cell therapy but with oral safety and simplicity.
4. UT-018 represents a new generation of regenerative nutraceuticals for epithelial and gut health.

## Acknowledgements

We thank Cology Biosciences Pvt. Ltd., an accredited and certified in vivo Contract Research Organization, for conducting the DSS-induced colitis study and histopathological evaluations. All studies were performed in accordance with the Institutional Animal Ethics Committee (IAEC) approval and complied with **CPCSEA** guidelines. Authors are affiliated with Utopia Therapeutics, developer of UT-018.

